## Supporting Information for "A drop dispenser for simplifying on-farm detection of foodborne pathogens"

**for**

**Supporting Information**

**Figure S1**. A comparison between the performance of some commercial pipettors versus our drop dispenser. All values are in µL.

**Figure S2**. A calibration curve to estimate the *E. coli* O157:H7 cell counts from $OD_{600}$ reads.

**Figure S3.** Quantification of the *E. coli* O157:H7 DNA extract using PicoGreen dye. A) Correlating the Lambda DNA concentration with fluorescence intensity. B) Estimating template DNA concentration.

**Figure S4.** Effects of various surface treatment procedures on the dispensing capacity of the drop dispenser. A) Importance of having a hydrophilic interior and hydrophobic exterior surface for the liquid holder of the drop dispenser on the dispensing performance. B) Performance of drop dispensers treated by plasma and PEG 400 after a while of storage. All values are in µL

**Figure S5.** Effect of resin types on the performance of the drop dispensers. All values are in µL.

**Figure S6a.** A comparison of the LAMP assay results (replicate 1) when using our drop dispenser versus a standard Eppendorf 20-200 µL pipettor. A yellow color indicates a positive result of the test. The template was STEC O157:H7 DNA extract at various dilutions.

**Figure S6b.** A comparison of the LAMP assay results (replicate 1) when using our drop dispenser versus a standard Eppendorf 20-200 µL pipettor. A yellow color indicates a positive result of the test. The template was *E. coli* O157:H7 DNA extract at various dilutions.

**Figure S6c.** A comparison of the LAMP assay results (replicate 1) when using our drop dispenser versus a standard Eppendorf 20-200 µL pipettor. A yellow color indicates a positive result of the test. The template was *E. coli* O157:H7 DNA extract at various dilutions.

**Figure S7.** A comparison of the whole-cell LAMP assay results when using our drop dispenser versus a standard Eppendorf 20-200 µL pipettor. A yellow color indicates a positive result of the test. The template was *E. coli* O157:H7 cells at various dilutions.

**Figure S8.** Results of LOD tests for whole-cell LAMP assays using *E. coli* O157:H7 cells at various dilutions. A yellow color indicates a positive result of the test. All liquid handling were performed using a standard Eppendorf 20-200 µL pipettor.

**Table S1.** LAMP primer set used to detect *E. coli* O157:H7 by targeting *stxI* gene (Wang et al., 2012).

**Table S2.** Bill of materials for colorimetric LAMP using drop dispensers (prices in USD).

**Design files.** The .stl files for the drop dispenser (1) plunger and (2) liquid holders are included

**Performance of commercial micro-pipettors vs. our devices**

| Micro-pipettor | Test 1 | Test 2 | Test 3 |
|---|---|---|---|
| Our device | 22.4±1.18 | 21.8±1.00 | 22.2±0.69 |
| Drummond Wiretrol® II (20 µL) (Cat. No. 5-000-2020) | 20.4±1.47 | 20.6±1.30 | 21.1±1.40 |
| Drummond Microcaps® (25 µL) (Cat. No. 1-000-0205) | 16.5±2.14 | 14.4±1.02 | 13.9±0.54 |
| Squeezing dropper of BD Veritor™ At-Home COVID-19 Test kit (Ref. No. 256094) | 27.2±2.31 | 30.7±2.83 | 46.3±2.85 |
| Corning® 20-200 µL Lambda™ Plus Single-channel Pipettor (A standard laboratory pipettor) (set at 20 µL) | 20.1±0.36 | - | - |

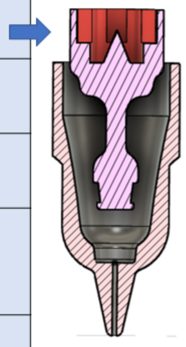

Figure S1. A comparison between the performance of some commercial pipettors versus our drop dispenser. All values are in µL.

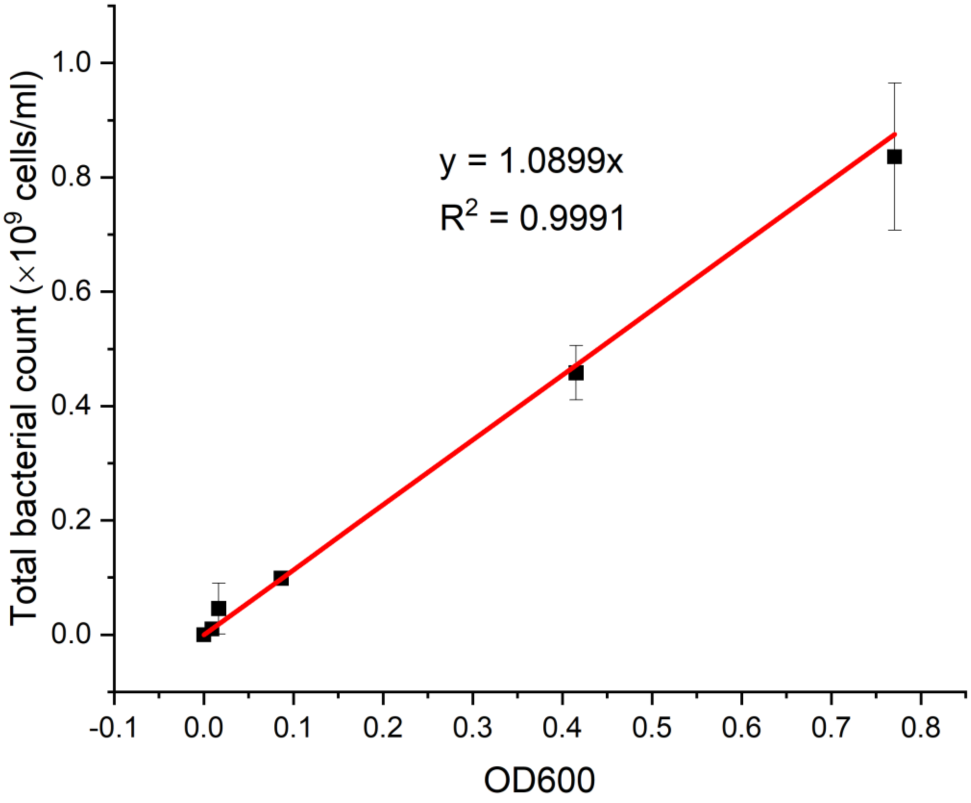

Figure S2. A calibration curve to estimate the *E. coli* O157:H7 cell counts from $OD_{600}$ reads.

### A) Correlating Lambda DNA concentration with fluorescence intensity

| | Fluorescence intensity | | | | |
|---|---|---|---|---|---|
| **Lambda DNA (ng/µl)** | **Replicate 1** | **Replicate 2** | **Replicate 3** | **Average** | **Standard deviation** |
| **2** | 112141.3 | 113443.9 | 107838 | 111141.1 | 2933.709 |
| **1** | 47476.48 | 47070.09 | 47197.58 | 47248.05 | 207.8426 |
| **0.5** | 25741.36 | 27928 | 28148.95 | 27272.77 | 1330.832 |
| **0.25** | 15550.63 | 15381.13 | 14793.06 | 15241.61 | 397.5854 |
| **0.125** | 10099.82 | 10537.89 | 10081.84 | 10239.85 | 258.2667 |
| **0.0625** | 6864.617 | 7631.577 | 7514.979 | 7337.058 | 413.2781 |
| **0.03125** | 5916.694 | 5966.507 | 6090.179 | 5991.127 | 89.32475 |
| **0.015625** | 5044.878 | 5250.53 | 4998.129 | 5097.846 | 134.2785 |
| **0** | 4455.722 | 6413.868 | 4801.635 | 5223.741 | 1045.091 |

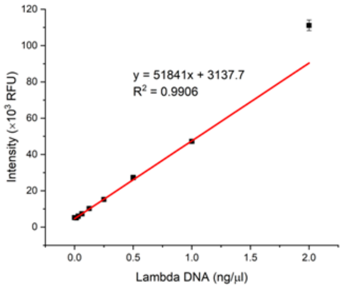

### B) Estimating template DNA concentration

#### Fluorescence intensities for template DNA dilutions

| | Fluorescence intensity | | |
|---|---|---|---|
| **Template DNA (*E coli* O157)** | **Replicate 1** | **Replicate 2** | **Replicate 3** |
| **DNA stock** | ? | ? | ? |
| **DNA dilution 1 (1:10)** | 114097.2 | 114599.4 | 150003 |
| **DNA dilution 2 (1:10)** | 112861.8 | 113175.7 | 137759.4 |
| **DNA dilution 3 (1:2)** | 75780.24 | 73776.85 | 74509.3 |
| **DNA dilution 5 (1:2)** | 39503.64 | 39683.61 | 39110.59 |
| **DNA dilution 5 (1:2)** | 22866.63 | 23960.5 | 25484.13 |
| **DNA dilution 6 (1:2)** | 13771.15 | 13121.44 | 13714.47 |

#### Estimated DNA concentration in ng/µl

| Template DNA predicted (ng/µl) | | | | |
|---|---|---|---|---|
| **Replicate 1** | **Replicate 2** | **Replicate 3** | **Average** | **Standard deviation** |
| | | | 23.74482 | |
| 2.14038 | 2.150069 | 2.832996 | 2.374482 | 0.397114 |
| 2.11655 | 2.122606 | 2.596819 | 2.278658 | 0.275552 |
| 1.401257 | 1.362612 | 1.37674 | 1.380203 | 0.019554 |
| 0.70149 | 0.704962 | 0.693908 | 0.70012 | 0.005653 |
| 0.380566 | 0.401667 | 0.431057 | 0.40443 | 0.025359 |
| 0.205117 | 0.192584 | 0.204023 | 0.200575 | 0.006942 |

Figure S3. Quantification of the *E. coli* O157:H7 DNA extract using PicoGreen dye. A) Correlating the Lambda DNA concentration with fluorescence intensity. B) Estimating template DNA concentration.

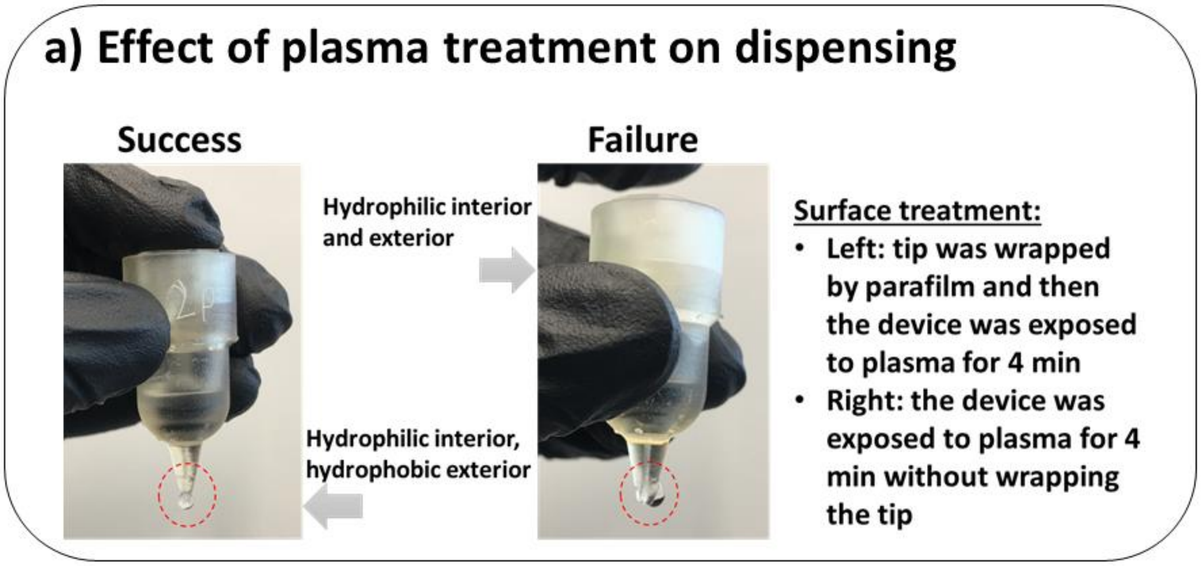

**b) Effect of surface treatment on dispensing capacity**

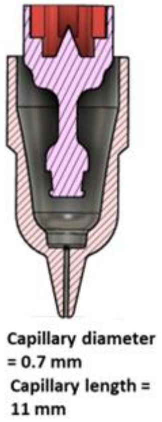

| | No PEG 400 | PEG 400 (for 24 hr) | After 35 days of PEG 400 treatment (for 24 hr) |
|---|---|---|---|
| No wrapped tip, No plasma | 16.8±1.72 | | |
| No wrapped tip, With plasma (for 2 min) | 20.8±2.78 | 24.1±1.67 | |
| Wrapped tip, With plasma (for 2 min) | 17.0±1.23 | 22.4±1.18 | 21.3±1.07 |
| Wrapped tip, With plasma (for 4 min) | 19.6±1.86 | | |
| Wrapped tip, After 10 days of plasma treatment (for 2 min) | 16.5±2.55 | | |

Figure S4. Effects of various surface treatment procedures on the dispensing capacity of the drop dispenser. A) Importance of having a hydrophilic interior and hydrophobic exterior surface for the liquid holder of the drop dispenser on the dispensing performance. B) Performance of drop dispensers treated by plasma and PEG 400 after a while of storage. All values are in µL.

**Effect of resin type on dispensing performance**

| Resin type | Device I | Device II | Device III | Device IV | Device V | Device VI |
|---|---|---|---|---|---|---|
| High Temp V2 | 22.4±1.18 | 21.8±1.00 | 22.2±0.69 | 21.9±1. 23 | 20.9±0.84 | 21.6±0.96 |
| Clear V4 | 21.8±1.22 | Failed* | Failed | Failed | Failed | 22.1±1.39 |
| Rigid 4000 V1 | Failed | Failed | Failed | Failed | Failed | Failed |

* The device failed due to uncontrolled dripping.

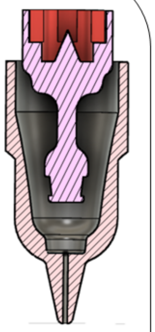

Figure S5. Effect of resin types on the performance of the drop dispensers. All values are in µL.

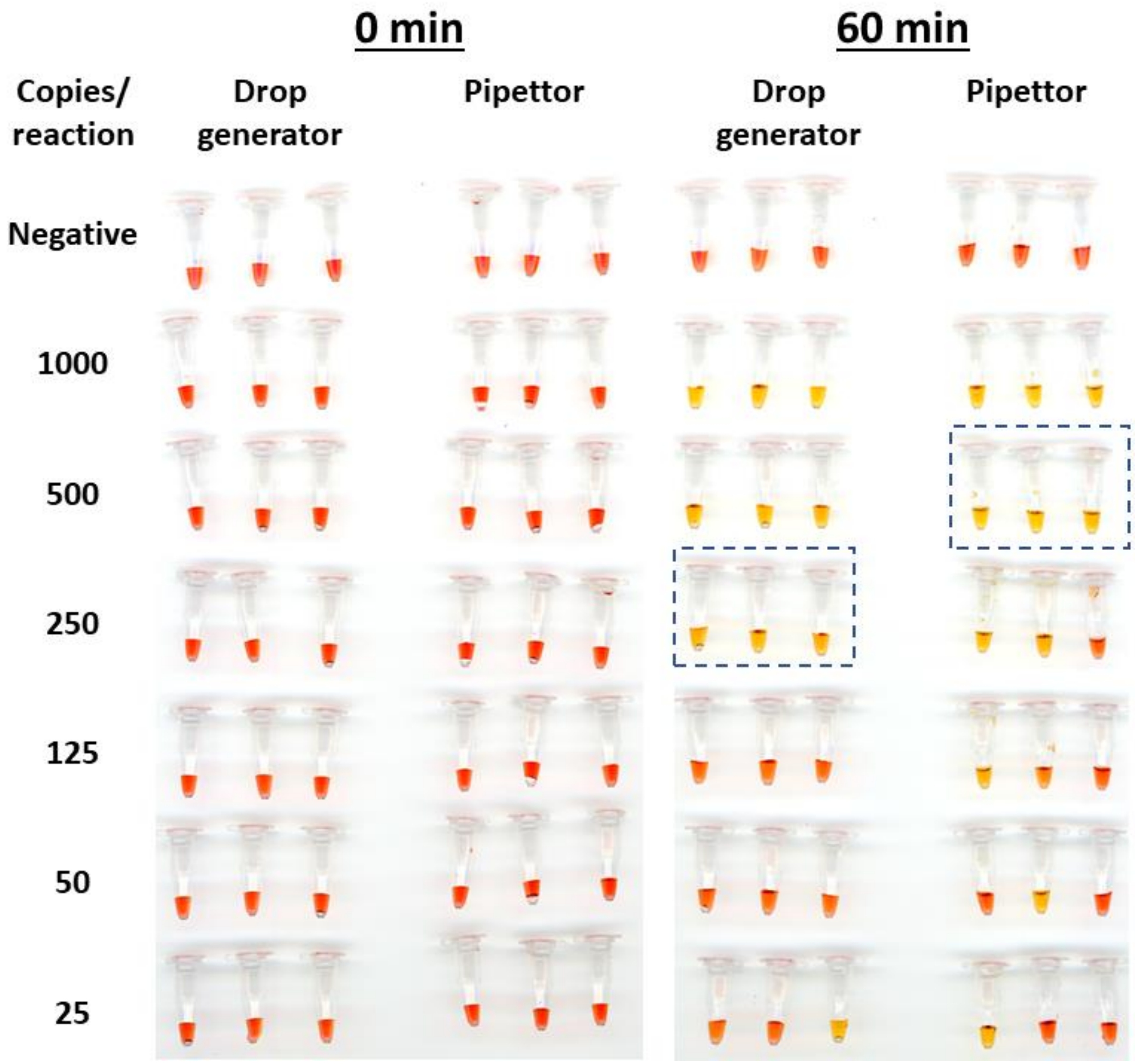

Figure S6a. A comparison of the LAMP assay results (replicate 1) when using our drop dispenser versus a standard Eppendorf 20-200 µL pipettor. A yellow color indicates a positive result of the test. The template was *E. coli* O157:H7 DNA extract at various dilutions.

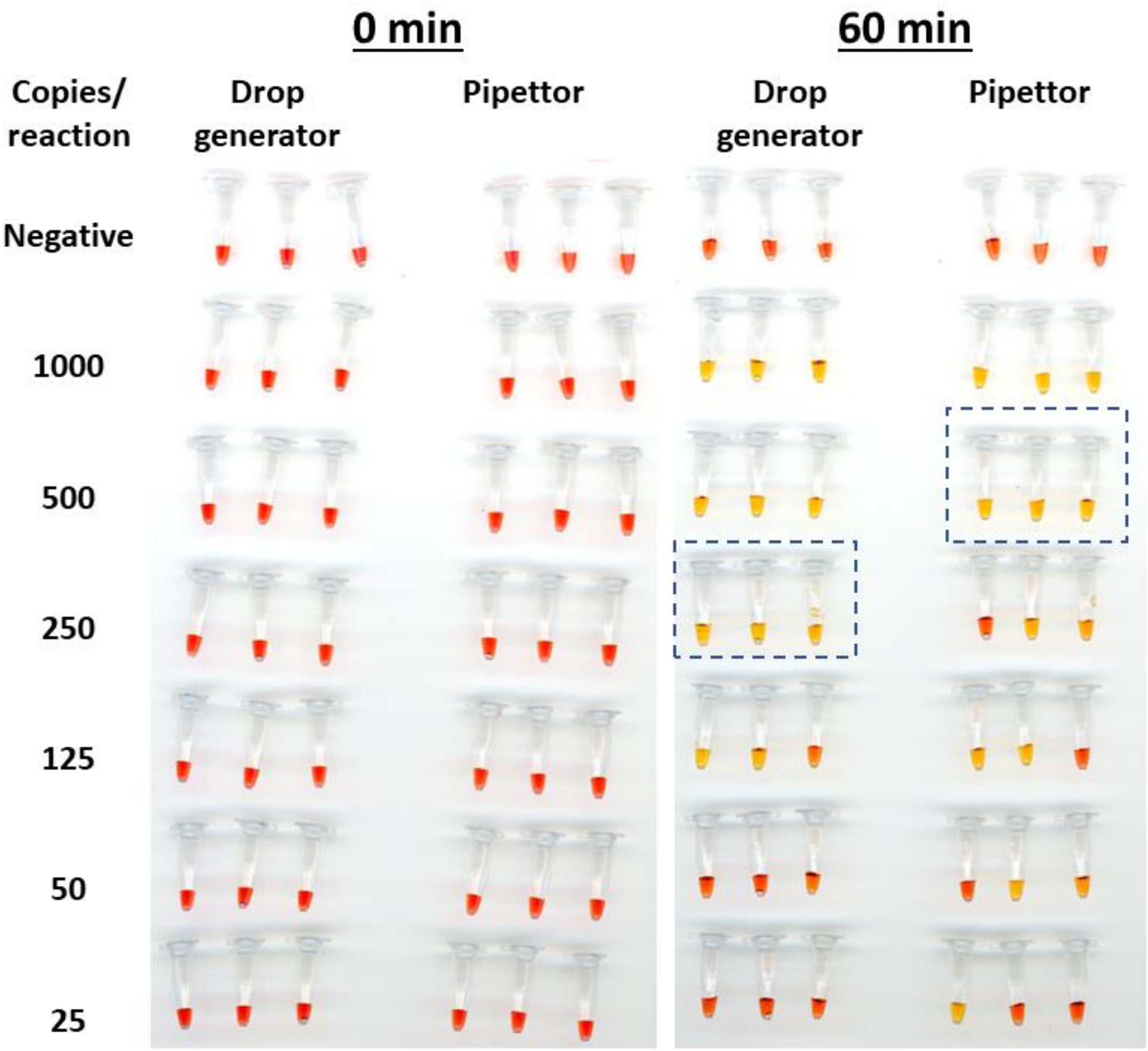

Figure S6b. A comparison of the LAMP assay results (replicate 1) when using our drop dispenser versus a standard Eppendorf 20-200 µL pipettor. A yellow color indicates a positive result of the test. The template was *E. coli* O157:H7 DNA extract at various dilutions.

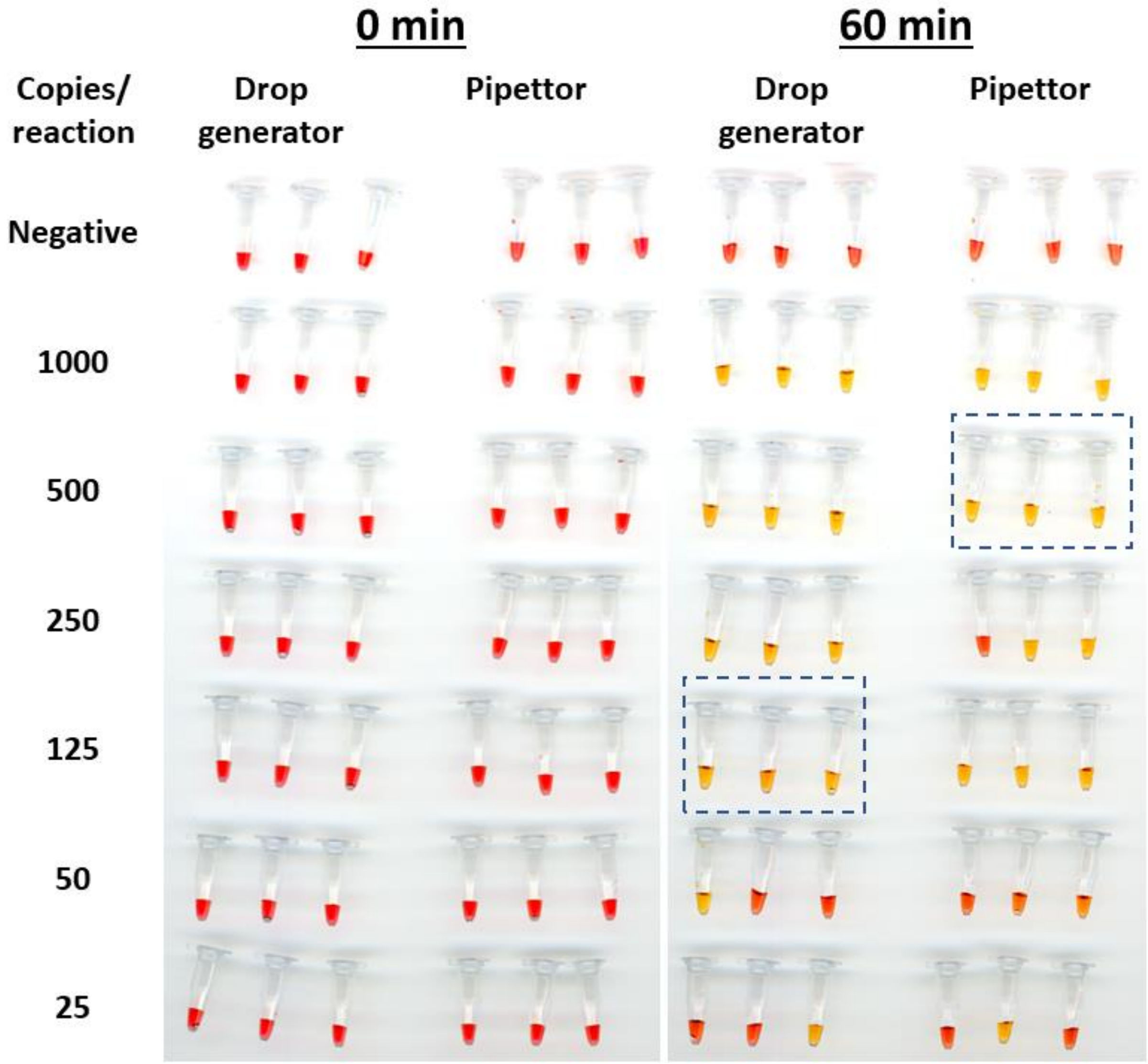

Figure S6c. A comparison of the LAMP assay results (replicate 1) when using our drop dispenser versus a standard Eppendorf 20-200 µL pipettor. A yellow color indicates a positive result of the test. The template was *E. coli* O157:H7 DNA extract at various dilutions.

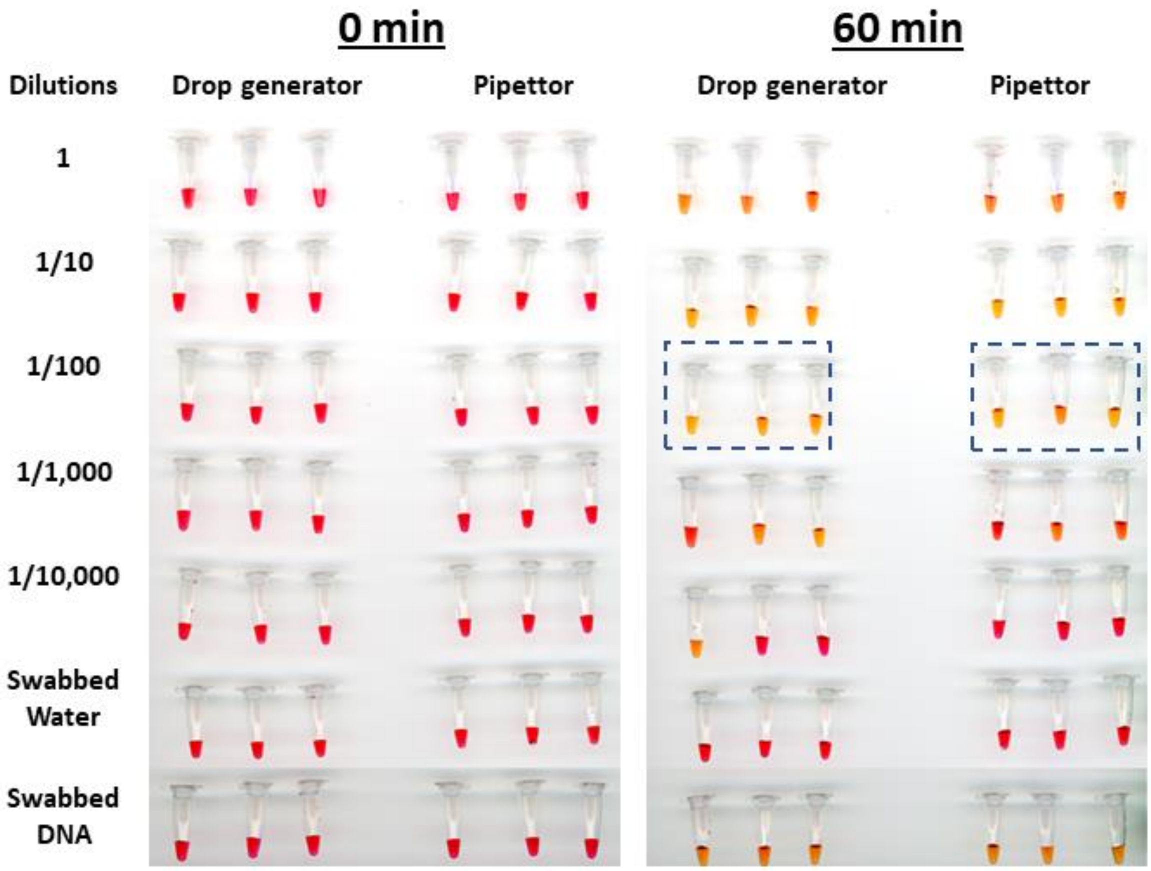

Figure S7. A comparison of the whole-cell LAMP assay results when using our drop dispenser versus a standard Eppendorf 20-200 µL pipettor. A yellow color indicates a positive result of the test. The template was *E. coli* O157:H7 cells at various dilutions.

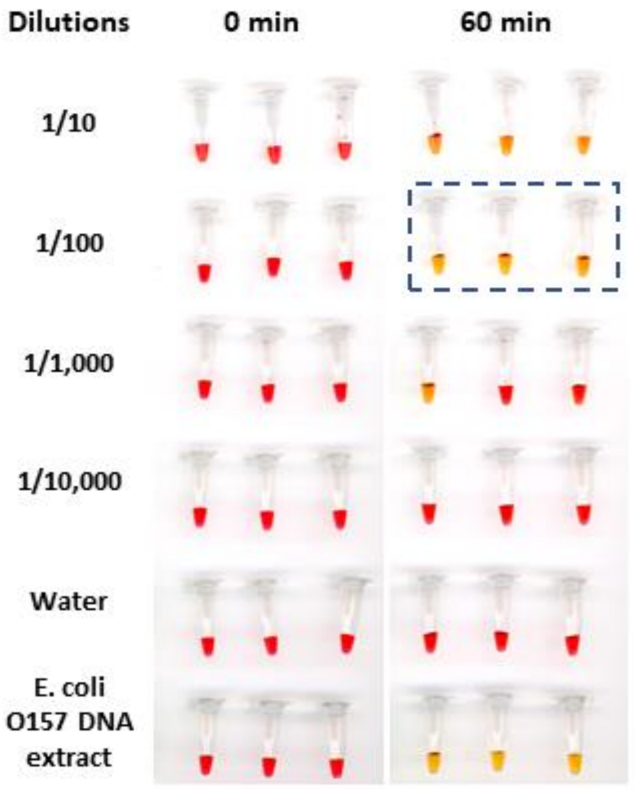

Figure S8. Results of LOD tests for whole-cell LAMP assays using *E. coli* O157:H7 cells at various dilutions. A yellow color indicates a positive result of the test. All liquid handling were performed using a standard Eppendorf 20-200 µL pipettor.

Table S1. LAMP primer set used to detect *E. coli* O157:H7 by targeting *stxI* gene

| Primer name | Sequence (5’-3’) |
|---|---|
| **EC.stx1-F3** | TGATTTTTCACATGTTACCTTTC |
| **EC.stx1-B3** | TAACATCGCTCTTGCCAC |
| **EC.stx1-FIP** | CCTGCAACACGCTGTAACGTCAGGTACAACAGCGGTTA |
| **EC.stx1-BIP** | AGTCGTACGGGGATGCAGATAGTGAGGTTCCACTATGC |
| **EC.stx1-LF** | GTATAGCTACTGTCACCAGACAATG |
| **EC.stx1-LB** | AAATCGCCATTCGTTGACTACT |

Table S2. Bill of materials for colorimetric LAMP using drop dispensers (prices in USD)

| **Item** | **Source** | **Unit price** | **Amount of use per test** | **Price per test** |
|---|---|---|---|---|
| **Droplet dispenser** | | | | |
| High Temp V2 resin | Formlabs, RS-F2-HTAM-02 | $199/1 L | 10 mL | $1.99 |
| O-ring | Helipal, Airy-Acc-Oring-2.5×6mm | $2.9/40 pieces | 2 pieces | $0.15 |
| **Sub-total** | | | | **$2.14 (55 %)** |
| **LAMP assay** | | | | |
| Magnesium Sulfate | Sigma-Aldrich, M2773 | $68.20/500 g | 0.0002 g | $0.00 |
| Potassium Chloride | Sigma-Aldrich, P9541 | $47.80/500 g | 0.0004 g | $0.00 |
| Antarctic Thermolabile UDG | New England Biolabs, M0372S | $80.00/100 U | 0.0875 U | $0.07 |
| dNTPs | Fisher Scientific, FERR0182 | $634.50/4 mL | 0.0014 mL | $0.22 |
| dUTP | Fisher Scientific, FERR0133 | $80.70/250 µL | 0.0875 µL | $0.03 |
| Phenol Red | Sigma-Aldrich, P3532 | $105.00/25 g | 0.0094 g | $0.04 |
| Betaine | Sigma-Aldrich, B0300-5VL | $103.00/7.5 mL | 0.0001 mL | $0.00 |
| EC.stx1 Primer mix | Life Technologies, N/A | $68.44/12500 reactions | 1 reaction | $0.01 |
| Warmstart *Bst* 2.0 DNA Polymerase | New England Biolabs, M0537M | $296.00 /0.067 mL | 0.0001 mL | $0.44 |
| **Sub-total** | | | | **$0.81 (21 %)** |
| **Sample collection** | | | | |
| BD BBL Dacron Polyester-Tipped Swabs | BD, 263000 | $189.5/500 swabs | 1 swab | $0.85 |
| Transparency film | Apollo, 617993 | $32.99/100 sheets | 0.3 sheet | $0.10 |
| **Sub-total** | | | | **$0.95 (24 %)** |
| **Total** | | | | **$3.9** |

Design File: Image below shows the the assembly and cross-sectional view of the design files.

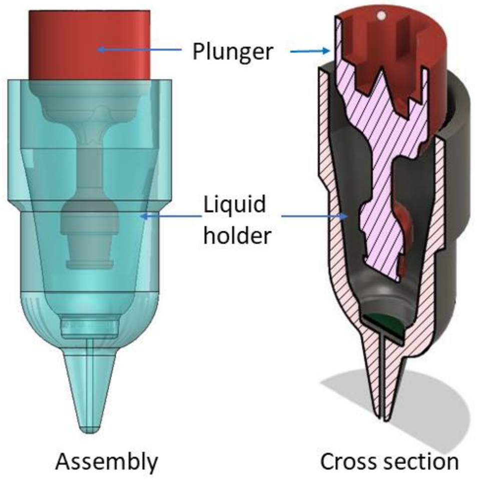
